## Supplemental Figures for "Integrating long-read RNA sequencing improves locus-specific quantification of transposable element expression"

### Supplementary Figures

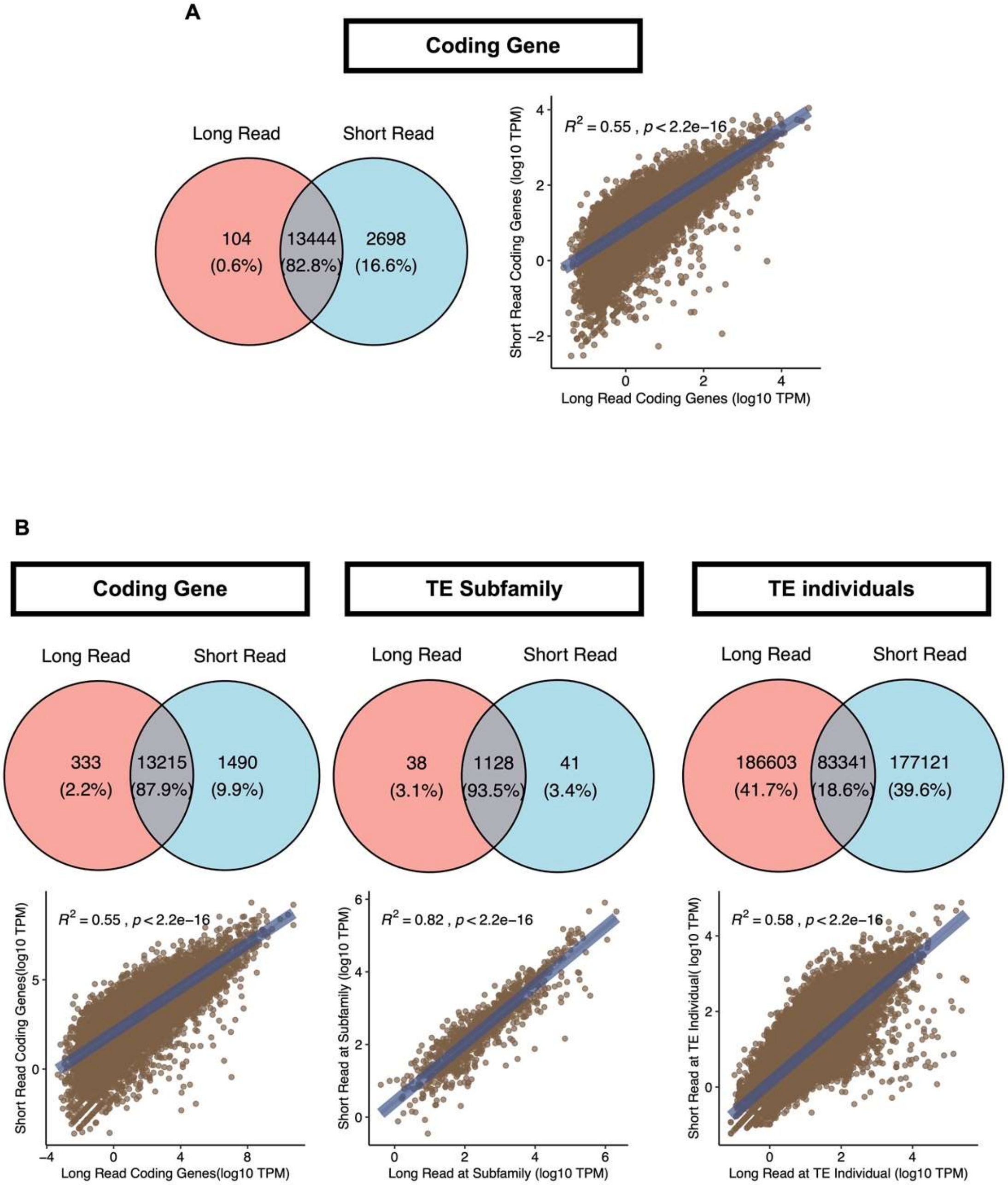


**Supplementary Figure 1: Comparison between short-read and long-read**
(A) Comparison of coding genes captured by short-read and long-read RNA-seq. Venn diagram showing the number of captured genes in two RNA-seq techniques. Correlation plot between TPM count for short-read and long-reads in the coding genes domain. (B) Comparison between downsampled short-read and long-read RNA-seq. The short-read sample is downsampled to match the number of bases mapped between short-read and long-read. Venn diagrams for downsampled short-read and long-read of TEs in subfamily and individual levels. Correlation plots between TPM count for downsampled short-read and long-reads in the same domains.

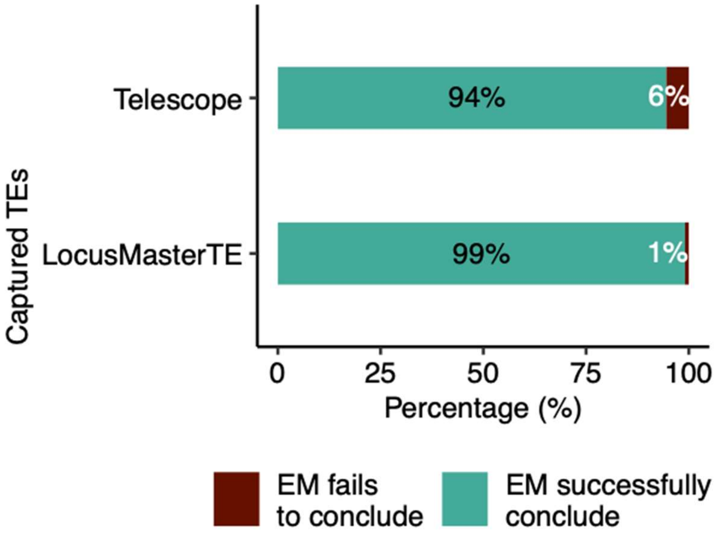


**Supplementary Figure 2: A comparison between Telescope and LocusMasterTE in the proportion of cases where EM successfully identifies the best hit**

Bar plot showing the proportion of cases that EM fails to determine a single genomic position that multi-mapped reads mapped to. Aided by long-read, LocusMasterTE reduced those cases to 1%.

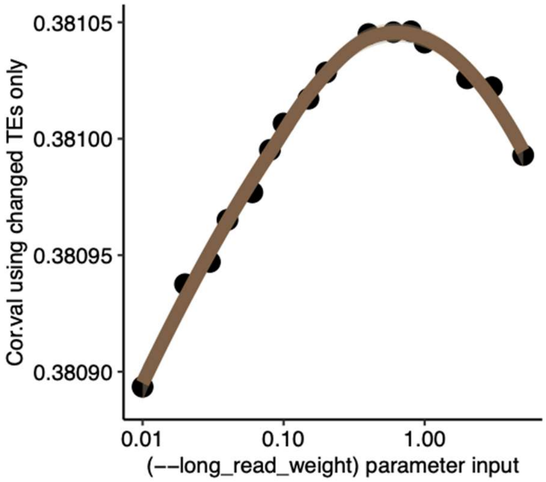


**Supplementary Figure 3: Impact of (--*long_read_weight*) parameter in LocusMasterTE**
TE expressions of the inhouse HCT116 sample with K562 long-read with varied (--*long_read_weight*) values and a series of correlation tests were conducted with the inhouse HCT116 long-read sample. Log TPM values were used for all TE expressions. For clear demonstration, only TEs with expression changed with (--*long_read_weight*) values are considered.

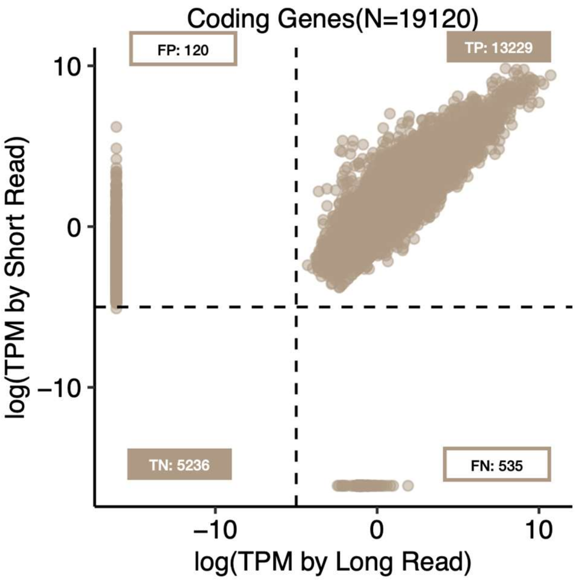


**Supplementary Figure 4: Simulated short-read for coding genes**
Correlation plot for 19120 coding genes between simulated short-read and long-read. This analysis is performed to check the success of the simulation. As expected, most of the coding genes are correctly simulated.

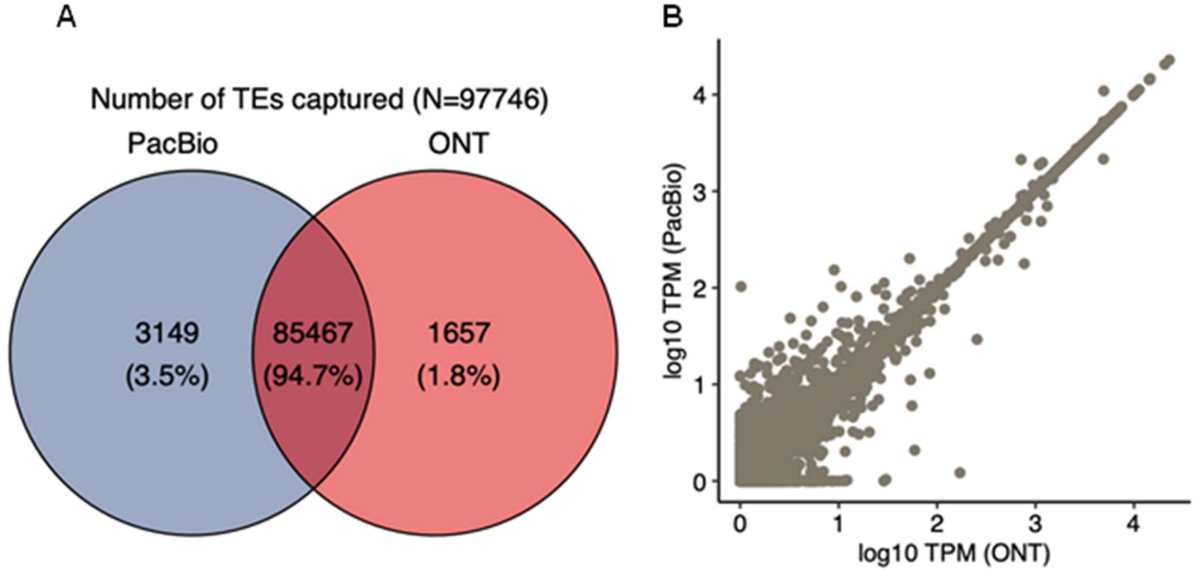


**Supplementary Figure 5: Quantified TE expression between Nanopore vs PacBio on simulated short-read**

(A) Venn diagram showing the number of TE individuals captured by LocusMasterTE using PacBio vs. ONT. (B) Correlation plot between quantified TEs by LocusMasterTE using PacBio as long-read input vs ONT as long-read input.

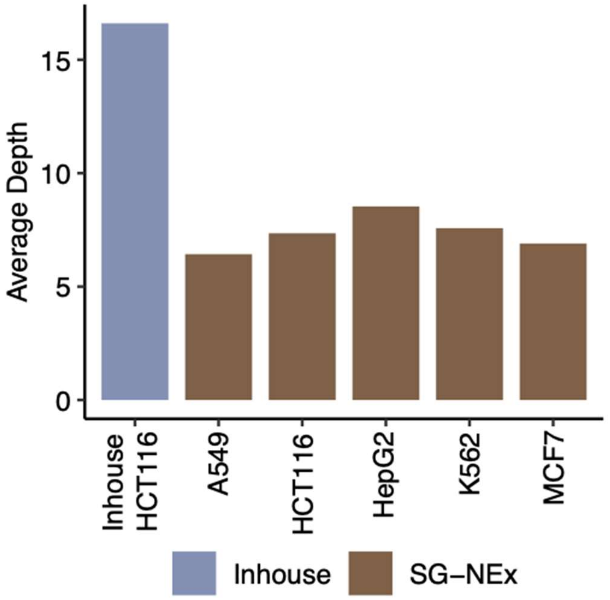


**Supplementary Figure 6: Average depth across all 6 samples**
Bar plot showing average read depth across all 6 samples.

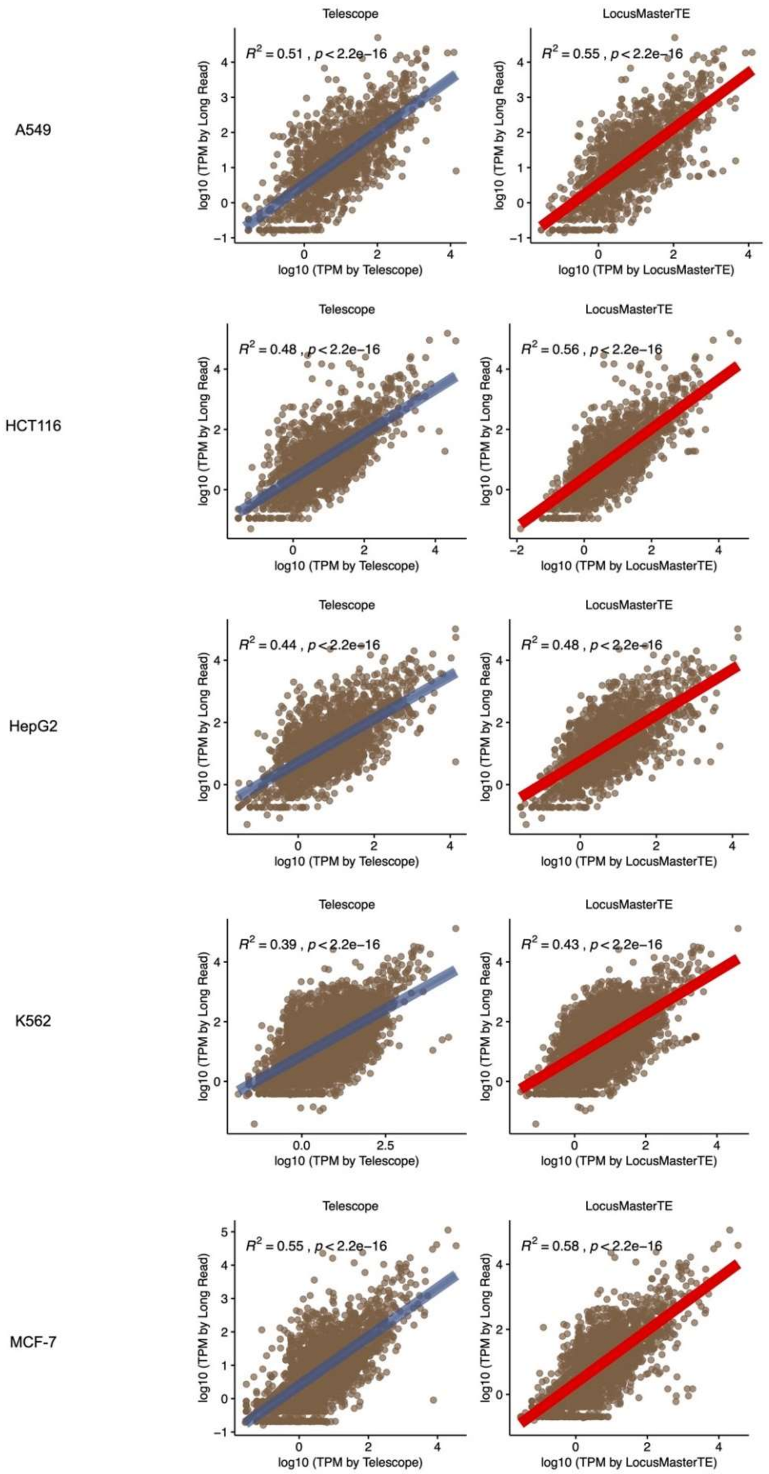


**Supplementary Figure 7: Correlation plot between long-read and short-read for SG-Nex**
Series of correlation plots between long-read and short-read for 5 SG-NEx cell lines. Telescope and LocusMasterTE quantify TEs in 5 cell line samples. Generated TE matrices are compared with the long-read result. Brown: TEs quantified higher in LocusMasterTE (TEs measured highly by LocusMasterTE). Blue: TEs are quantified higher in Telescope [8] (TEs are measured highly by Telescope).

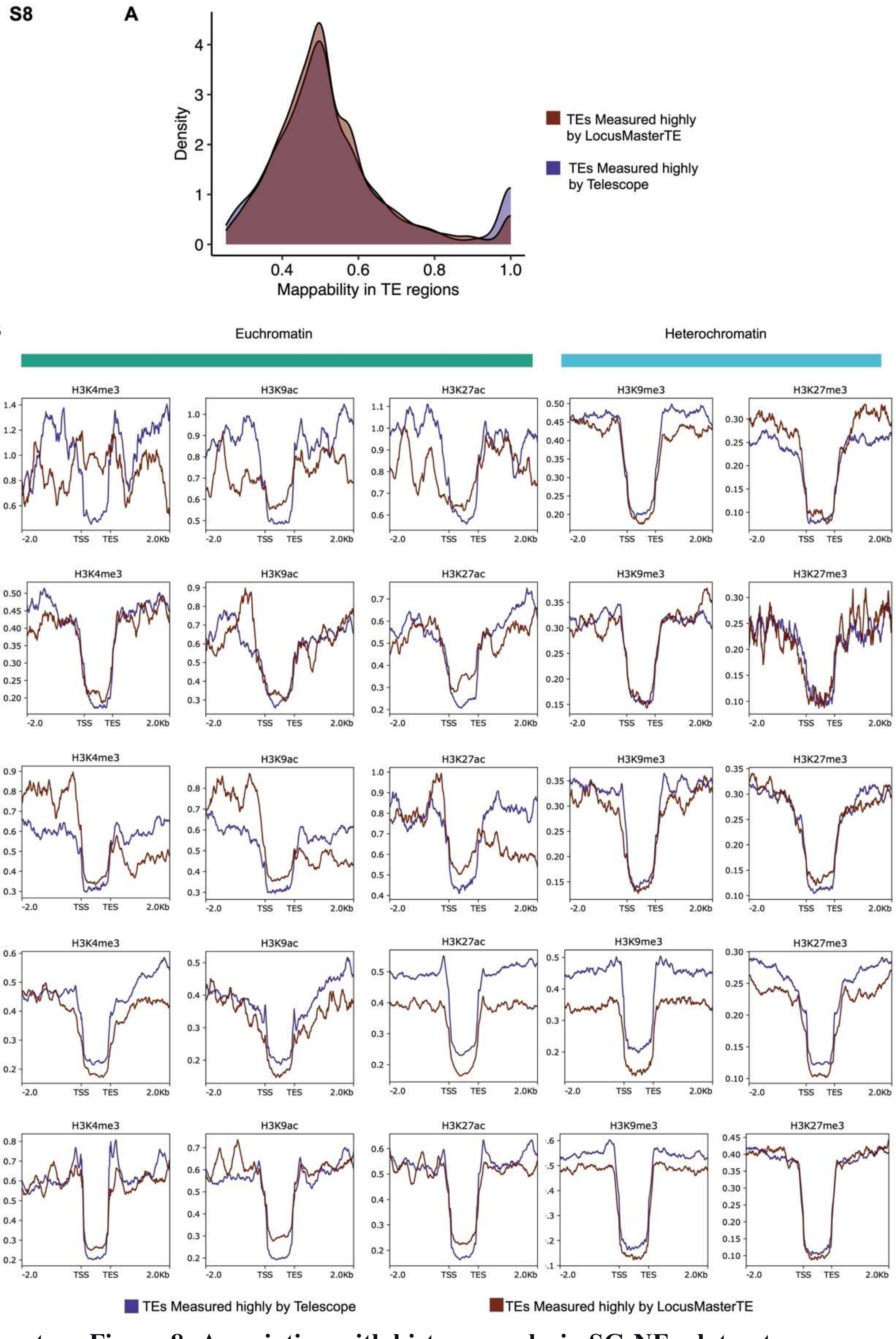


**Supplementary Figure 8: Association with histone marks in SG-NEx dataset.**
(A) Distribution graph of average mappability in two groups of TE regions. Mappability > 0.25 was selected. Brown: TEs quantified higher in LocusMasterTE (TEs measured highly by LocusMasterTE). Blue: TEs are quantified higher in Telescope [8] (TEs are measured highly by Telescope). (B)Three active chromatins coverage of 5 SG-NEx samples for two TE groups: TEs quantified higher in LocusMasterTE (TEs measured highly by LocusMasterTE) and TEs quantified higher in Telescope (TEs measured highly by Telescope). Two repressive chromatins coverage of 5 SG-NEx samples for two TE groups, same as above.
